# Correlative 4D-STEM Ptychography and EELS for Cryogenic Single-Particle Analysis

**DOI:** 10.1101/2025.10.21.683550

**Authors:** Xiaopeng Wu, Yu Lei, Peng Wang

## Abstract

Cryogenic electron microscopy has enabled atomic resolution structural determination of biological macromolecules. However, phase contrast imaging alone does not reveal elemental composition or other valuable chemical information. Electron energy loss spectroscopy (EELS) provides rich vibrational and chemical details, integrating it with high-resolution structural imaging for correlative analysis of biological specimens remains a major challenge, as conventional acquisition modes are often mutually exclusive. Here we propose two optical geometries that enable correlative 4D-STEM with EELS: (1) a sequential acquisition with a post-specimen beam deflector, and (2) a simultaneous acquisition with a hollow detector. We compared the two setups for biological specimens, and focused on the hollow detector geometry which allows simultaneous acquisition of diffraction patterns and EELS spectra in a single dose budget. Using an apoferritin dataset, we demonstrate sub-nanometre 3D resolution in single particle analysis, even with a hollow angle as of 55% of the convergence angle and approximately 30% of the total dose allocated to EELS. Finally, we propose a workflow that integrates ptychography with EELS for correlative, energy-resolved 3D mapping of biological macromolecules, paving the way toward unified structural, chemical, and vibrational analysis of biological specimens.

## 1. Introduction

Cryogenic electron microscopy (cryo-EM) has emerged as a powerful tool for determine high-resolution structure of biological macromolecules. This technique typically involves flash-freezing a purified sample in a thin layer of vitreous ice, which preserves the molecules in their near-native, hydrated conformation. A transmission electron microscope (TEM) is then used to capture a multitude of two-dimensional projection images of the particles, that are randomly oriented within the ice layer. Cryo-EM has been demonstrated to determine near atomic resolution structures of proteins^1,2^, however due to the lack of elemental mapping, the interpretation of experimental densities into the corresponding atomic structure often relies on substantial prior information of the macromolecular complex. For components of a structure that have less prior information readily available, such as bound cofactors, lipids, substrates, inhibitors, post-translational modifications as well as metals and other ions, this leads to more uncertainty in the final resolved structures^3^.

Alternative imaging methods can offer complementary information. Fluorescence microscopy provides molecular specificity, but it does not resolve ultrastructural details. Fluorescence can be combined with EM through correlative approaches, but suffer from issues such as loss of labels during sample preparation or accurately aligning datasets that have different resolution levels. It remains challenging to directly links structure with elemental composition in the cryo-EM environment itself.

While both energy-dispersive X-ray spectroscopy (EDS) and electron energy-loss spectroscopy (EELS) can, in principle, provide chemical information in the electron microscope, EELS offers several advantages over EDS in the case of biological specimens^3^. EELS is highly sensitive to light elements (H, C, N, O, S and P), of which biomacromolecules are primarily composed of^4,5^. This allows using a range of endogenous elements as label-free intrinsic marker for different biological components. For example, the phosphorous L_2,3_ edge at 132 eV signals the location of phosphate groups in nuclei acids or phosphorylated proteins^6-9^, and the sulphur L_2,3_ edge at 165 eV can be used an indicator for sulphur-containing amino acids^10^. By analysing the K-edge of carbon, nitrogen and oxygen, the ratios of these major constituents of organic molecules can be quantified^11^. In addition, EELS is sensitive enough to detect and map essential metal ions that function as cofactors or signalling molecules, such as calcium or iron, often at near-trace concentrations^12-14^. This ability to visualise the distribution of important elements is crucial for understanding cellular physiology and pathology.

Crucially, EELS is also more dose-efficient for thin cryo-specimens, making it compatible with the stringent requirements of cryo-EM. While both techniques require a very large effective dose to achieve single-atom resolution, EELS detects the ionisation events directly instead of relying on subsequent X-ray radiation, which has limited yield for light elements. The directions of inelastically scattered electrons are mostly concentrated around the direction of the incident electron beam, meaning the collection efficiency at the spectrometer can be very high in comparison to characteristic X-rays that are emitted isotopically in all directions.

Moreover, EELS can provide more chemical analysis beyond merely elemental composition, where fine structures in the low-loss and near edge region can reveal important details about the local composition, bonding, and oxidation states of the detected atom^4^, this would allow distinguishing between the same elemental species that serves different biological roles. An emerging capability of EELS provides access to vibrational modes of molecules (vibEELS) when combined with monochromated beams, providing information about local bonding, secondary and tertiary structure, hydrogen bonding networks, ligand coordination, and conformational dynamics. Spatial resolutions of order ∼10 nm with meV energy resolution have been demonstrated, directly resolving bond-specific signatures such as C such as C–H, C=O, N-H, or C–F functional groups in biological structures^15^.

When integrated with scanning transmission electron microscopy (STEM), EELS can be operated in parallel with a high-angle annular dark field (HAADF) imaging mode, where elastically scattered electrons can be detected at high angle to produce a Z-contrast images that are sensitive to heavy elements, giving access to a comprehensive understanding of the sample’s structure and composition^16,17^. The combination of STEM-HAADF and EELS has been successfully applied to a wide range of materials^18-21^ in physical sciences, however, this imaging mode has some intrinsic limitations when it comes to imaging biological specimens. As the intensity of high-angle scattered electrons is approximately proportional to the square of the atomic number (Z^2^), STEM-HAADF is less sensitive to light elements, and has a very low dose efficiency as only electrons that have been scattered at very high angles would contribute to the image contrast, this severely limits its application to beam sensitive materials, such as biological specimens.

Recent developments in high-speed pixelated detectors have enabled fast acquisition of 4D-STEM dataset^22-24^, which contains the full diffraction image instead of only an integrated signal at each probe positions, this allows for more flexible postprocessing, give access to various imaging modes virtually, such as bright field (BF), annular dark field (ADF), HAADF and centre of mass (CoM) imaging, as well as electron ptychography. Ptychography is a computational technique that would allow dose-efficient recovery of both the amplitude and phase of the specimen via a reconstruction algorithm. Electron ptychography has been demonstrated experimentally to achieve spatial resolution beyond the conventional diffraction limit and poses excellent dose efficiency and phase sensitivity to both light and heavy elements, making it well suited to biological applications^25-28^.

Conventional electron ptychography requires the full diffraction patterns for phase reconstruction, which is naturally incompatible with EELS acquisition as in this case all the signals would be captured by the pixelated detector, blocking any inelastically scattered electrons from entering the spectrometer. Two practical solutions could be explored to address this issue. First, a beam-deflecting scheme uses a post-specimen deflector to alternately steer the scattered electrons to the conventional pixelated detector and to the EELS spectrometer either between pixels or frames, preserving complete diffraction patterns for ptychography while acquiring spectra sequentially at matched scan positions (Fig. 1, Setup (a)); this retains phase information but is not truly simultaneous and is more susceptible to drift. Second, a new hollow detector configuration was proposed by Song et al^29^, where a hollow opening on the pixelated detector is designed to allow for electrons in low-angle regions of the BF disk to pass through to the EELS spectrometer (Fig. 1, Setup (b)). This allows simultaneous phase and Z-contrast imaging along with spectroscopic analysis. The hollow ptychographic imaging mode has been demonstrated on a monolayer MoS_2_ sample with sub-ångström resolution and robust phase reconstruction, establishing the viability of the technique.

**Figure 1:**
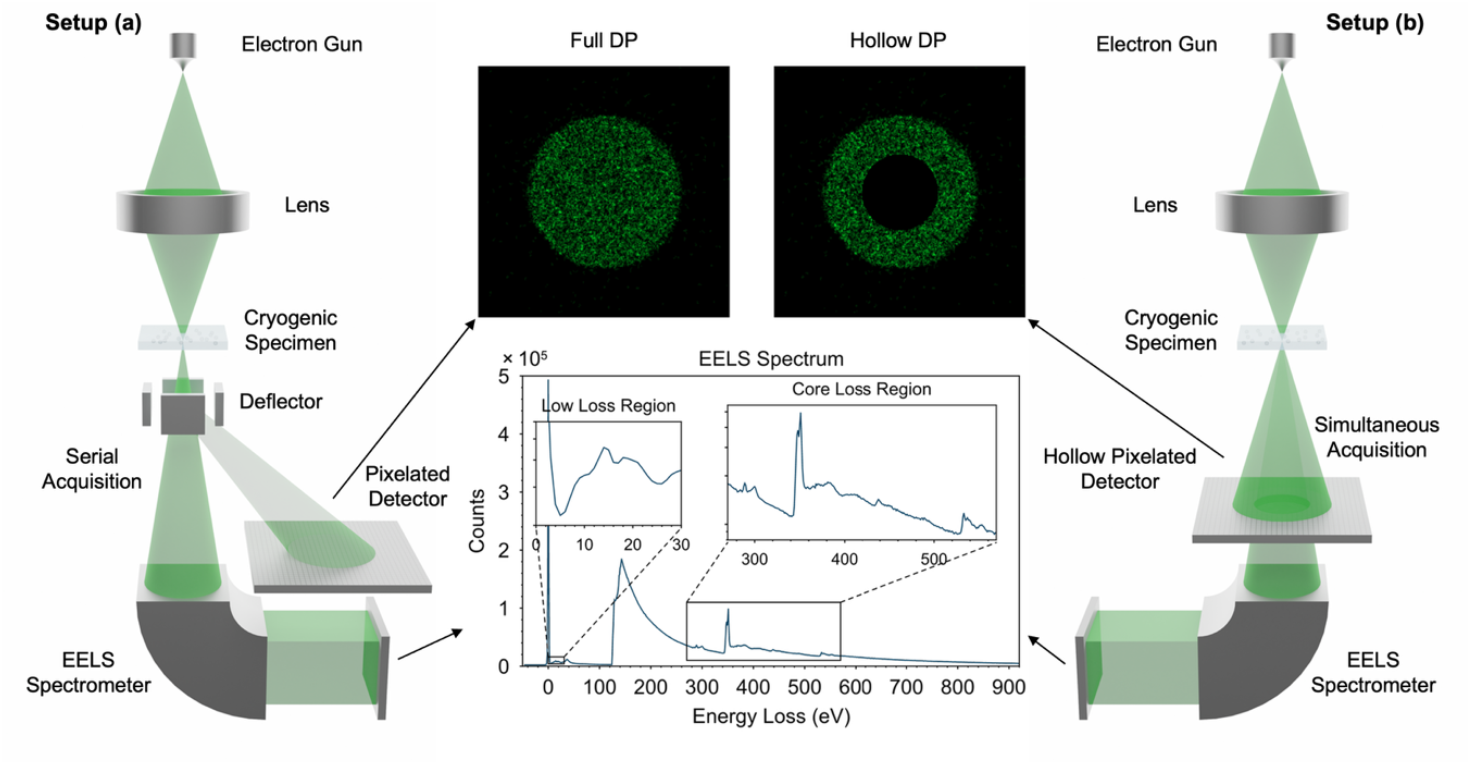
Schematic of experimental setups for correlative 4D-STEM with EELS. (a) Post-specimen beam-deflecting configuration: a post-specimen deflector alternates scattered electrons between a pixelated detector and the EELS spectrometer, acquiring full diffraction patterns and EELS spectra sequentially. (b) Hollow-detector configuration: a pixelated detector with a central opening pass through low-angle scattered electrons to the EELS spectrometer while recording the rest of the diffraction pattern, enabling simultaneous, co-registered structural and spectral acquisition. Illustrative EELS spectrum adopted from the EELS.info atlas^48^.

In this paper we present two experimental setups that enable electron ptychography with correlative EELS acquisition. We focused on the performance of hollow ptychography based low-dose 4D-STEM datasets of proteins under cryogenic conditions. We evaluated the reconstruction quality at various hollow semi-angles (HSA) and show that sub-nanometre 3D resolution can still be achieved even after partitioning a significant percentage of electron dose to EELS. We propose a novel workflow for data collection and processing for hollow cryogenic electron ptychography with EELS. Our results show that hollow ptychography is a compelling alternative to STEM-HAADF at enabling simultaneous acquisition of high-contrast structural and chemical information from biological specimens.

## 2 Results

### 2D Hollow Ptychographic Reconstruction of Proteins

Apoferritin, the hollow spherical protein shell of the ubiquitous iron-storage protein ferritin, is a highly symmetrical and stable nanocage with a diameter of roughly 12 nm. These properties helped to establish apoferritin as a standard cryo-EM reference particle. In this study, we reuse the cryogenic apoferritin 4D-STEM datasets collected by Küçükoğlu et al^28^ to performance hollow ptychographic reconstructions, where the central region of the diffraction disk is progressively masked to simulate the increasingly large HSAs as percentage of the convergence semi-angle (CSA), α. All the ptychographic reconstructions obtained in this work were based on a batched stochastic gradient descent algorithm, modified from Py4DSTEM^30^ to enable hollow reconstruction. Further details of the reconstruction method can be found in Supplementary Information Note 2.

Fig. 2a–d show diffraction patterns masked at 0α, 0.55α, 0.75α, and 0.95α, with their corresponding phase reconstructions in Fig. 2e-h. Reconstructions remain visually comparable to the full-pattern case up to 0.75α. At 0.95α, both the contrast and sharpness of the apoferritin particles are largely degraded, due to increased dose partitioning to the EELS spectrometer. Magnified images in Fig. 2i-l reveal that only coarse particle outlines are visible with high-frequency details lost at HSA beyond 0.75α. This observation is consistent with line profiles extracted across the particles as shown in Supplementary Fig. 2.

**Figure 2:**
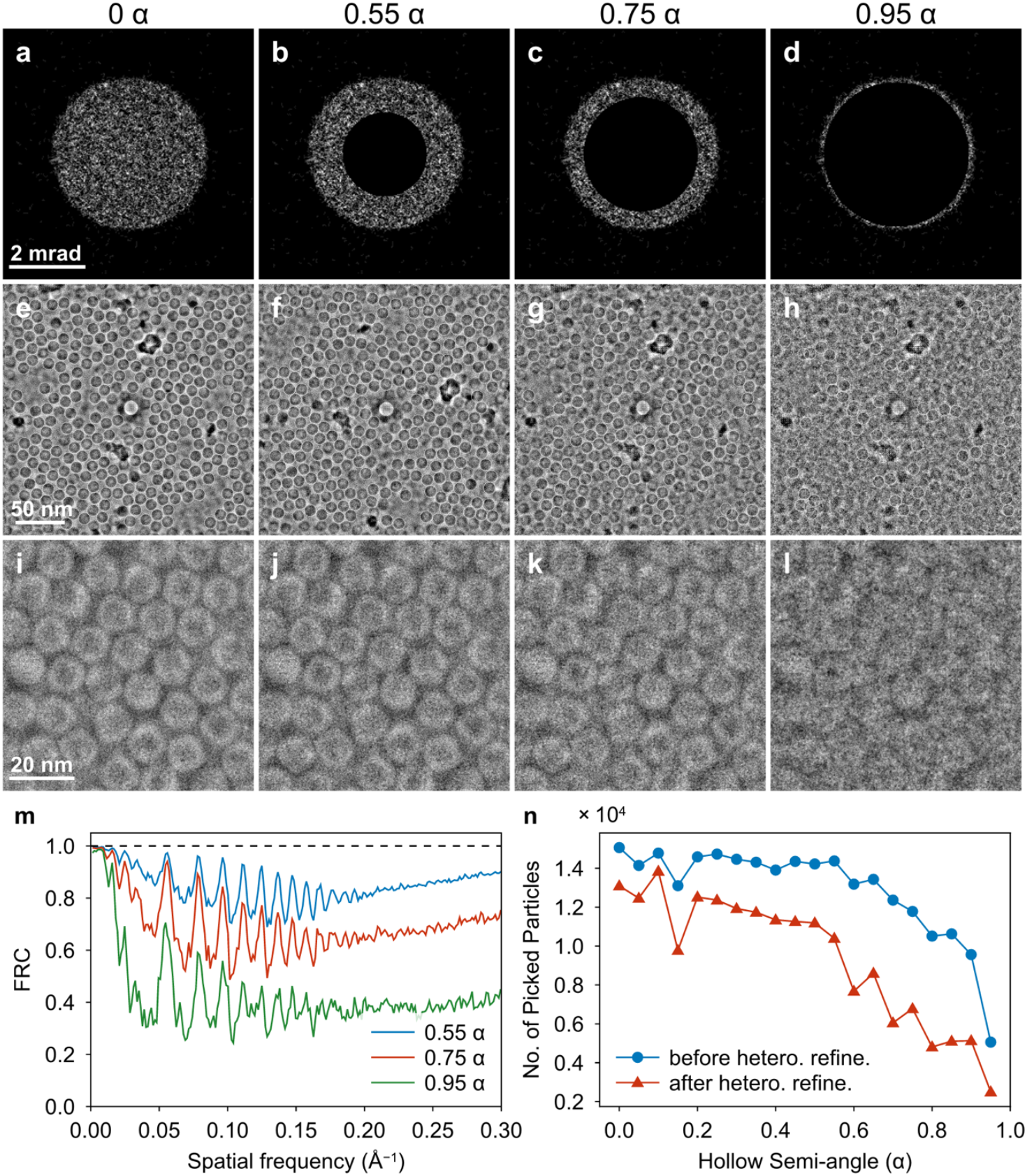
Reconstruction quality of apoferritin under varying hollow semi-angle (HSA) in hollow electron ptychography. **a**-**d** Diffraction patterns with different HSA values of 0, 0.55, 0.75, and 0.95 times the probe CSA. **e**-**h** Reconstructed phase images from the corresponding HSAs with picked particles highlighted with white circles, the number of picked particles are 260, 221, 125 and 47 respectively. **i**-**l** zoomed in view of the reconstructed apoferritin particles. **m** FRC curves of the reconstructions with different HSAs compared to the reconstruction from the full diffraction patterns (HSA = 0). **n t**otal number of particles used for single particle analysis from 90 individual micrographs reconstructed at different HSAs.

To further examine how HSA influences the information transfer across spatial frequencies, Fourier ring correlations (FRC) analyses (Fig. 2m) were performed relative to the image reconstructed with full diffraction patterns (Fig. 2e)^31^. At 0.55α and 0.75α, the FRC curves remain above 0.8 and 0.5, respectively, across most spatial frequencies, whereas at 0.95α the correlation drops below 0.5 except at the lowest frequencies.

The extent of reconstruction degradation at higher HSAs also affects the efficiency of particle picking, one of the first steps in cryo-EM data analysis performed using CryoSPARC^32^, in which individual particles of interest are identified and selected. As image quality declines, particle identification becomes increasingly difficult. Representative images with accepted picks are shown in Fig. 2e-h; the number of picks per image decreased from 260 at 0α to 47 at 0.95α. Total counts across 90 reconstruction images are summarised in Fig. 2n 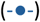. At HSA up to 0.75α, particle numbers remain at 80% of the count at 0α, then decline sharply to 50% at 0.95α (Supplementary Fig. 4). These trends are consistent with the significant loss of contrast and structural detail, which is largely preserved up to 0.75α but is substantially compromised at 0.95α and above.

2D classification showed the same HSA-dependent degradation (Supplementary Fig. 5). At higher HSAs, both the number and quality of usable classes declined substantially. Class averages became progressively noisier, with reduced contrast and loss of fine structural features, consistent with diminished high-frequency information in the underlying ptychographic reconstructions.

Furthermore, the number of the particles that will contribute to the final 3D reconstructions after the heterogeneous refinement was also assessed (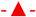 in Fig. 2n). During this refinement, the particle stack was separated into 2 subsets based on the structural variability exhibit in the dataset, the class with worse resolution was discarded, result in less available particle for further refinement. Across all HSAs, there was a substantial reduction in particle numbers compared with those picked prior to refinement. As shown in Supplementary Fig. 6, the percentage of the selected particles decrease with increasing HSA, suggesting that more particles fail the refinement at larger HSAs due to the degradation of reconstruction quality, making it harder for the particle to be assigned to a reference 3D class. This results in an additional loss of usable particles beyond the initial reduction during particle picking. Consequently, fewer particles enter *ab initio* reconstruction and refinement^32^, leading to reduced 3D resolution, as we will discuss in the next section.

### 3D Single Particle Reconstruction of Apoferritin

All *ab initio* reconstructions and subsequent refinements were performed in cryoSPARC^32^, which further methodological details are provided in Supplementary Information Note 3. Density maps reconstructed at HSAs of 0α, 0.55α, 0.75α, and 0.95α are shown in Fig. 3a-d, with their corresponding cross-sectional views as shown in Fig. 3e-h. At 0.5α, as at 0α, the reconstructed volume aligns well with the reference atomic model derived from atomic-resolution cryo-EM densities (PDB ID: 8RQB^28^). 3D resolution of the reconstructed volumes are evaluated using the gold standard Fourier shell correlation (gsFSC) with a 0.143 cutoff threshold (Supplementary Fig. 7)^33^, the resolutions at each HSAs are shown in Fig. 3m (-×-), which shows that the resolution at 0.55α is 6.92 Å, comparable to 6.64 Å at 0 α. By contrast, the resolution deteriorates dramatically to 8.34 Å, and 8.60 Å at HSAs of 0.75α, and 0.95α, respectively, preventing recovery of the correct secondary structure of apoferritin and are therefore not interpretable.

**Figure 3:**
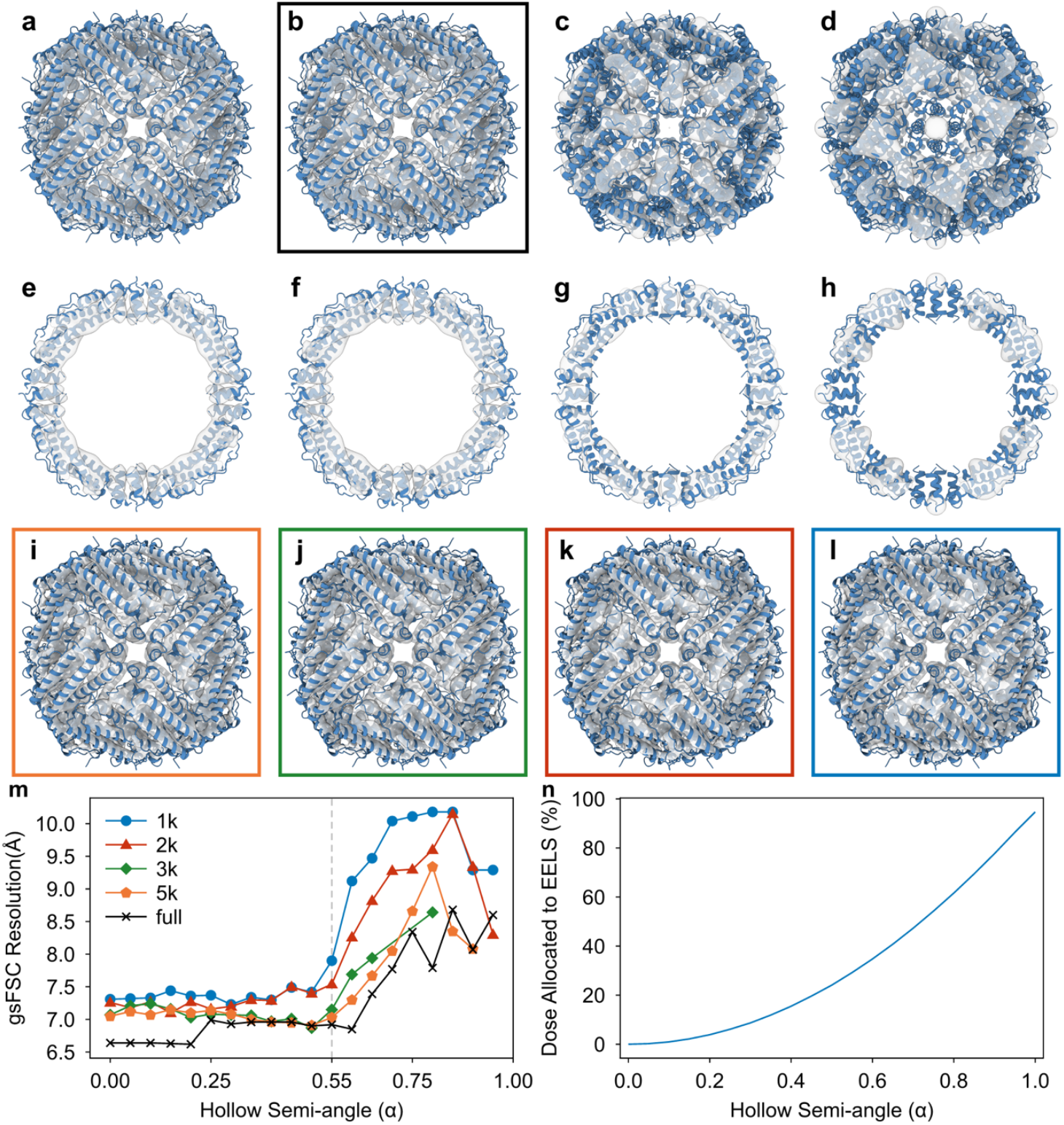
Three-dimensional reconstructions of apoferritin. **a**-**d** Density maps reconstructed from hollow ptychographic datasets with HSA values of 0, 0.55, 0.75, and 0.95 times the probe CSA, α, overlaid with the fitted atomic model (PDB ID: 8RQB). **e**-**h** Corresponding cross-sectional views. **i**-**l** Maps refined from 1,000, 2,000, 3,000, and 5,000 particles at HSA = 0.55α. **m** gsFSC (0.143 criterion) for reconstructions using 1,000, 2,000, 3,000, 5,000, and all available particles at each HSA. **n** Fraction of dose allocated to EELS versus HSA. Note: particle counts denote those used for the final local refinement and were randomly sampled from the best 3D class. For the “full” and 5,000-particle conditions, all available particles from the dataset were used for *ab initio* reconstruction; for smaller reconstructions, 5,000 particles were used.

Note that this evaluation was carried out using the same number of 4D datasets across all HSAs. As the HSA increases, both the quality and quantity of successful particle picks decrease. However, within the single-particle framework, the reduced quantity can in principle be compensated by acquiring additional 4D datasets, which may further improve the 3D reconstruction resolution.

To isolate the effect of 2D image quality variation (increasing HSA) on 3D reconstruction, we controlled particle numbers (quantity) at each stage. For every HSA, 5,000 particles were randomly drawn from the best heterogeneous refinement class and used to locally refine the structure. In parallel, *ab initio* reconstructions were performed using exactly 5,000 randomly selected particles to avoid bias from larger initial particle sets. From the 5,000-particle *ab initio* models, subsets of 1,000, 2,000, and 3,000 particles were randomly sampled from the best heterogeneous class for local refinement to assess resolution scaling with particle count under matched image quality (Supplementary Fig. 8). The resulting gsFSC resolutions are shown in Fig. 3m (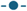 1,000, 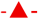 2,000, 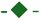 3,000, 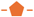 5,000, individual FSC curves can be found in Supplementary Fig. 9).

The impact of increasing HSA on achievable resolution depends strongly on particle count. The subsampled sets while never reaches ∼6.6 Å even at 0α comparing to the full set reconstruction show earlier inflection points: ≈0.5α (5,000 particles), ≈0.45α (3,000 particles), ≈0.30α (2,000 and 1,000 particles). Thus, indicating higher particle numbers can help mitigate the HSA-induced resolution drop likely by averaging noise. Using the reconstructions at 0.55α as an example, using 1,000 particles yielded 7.90 Å resolution, this improved to 7.52 Å with 2,000 particles and 7.15 Å with 3,000 particles; using 5,000 particles showed only minor degradation relative to the full dataset (6.92 Å). The density maps of these reconstructions are shown in Fig. 3b and Fig. 3i-l. These results indicate that increasing the particle count improves reconstruction quality, suggesting the high-frequency information is still partially retained in the ptychographic reconstructions at moderate HSAs but would require sufficient dose to recover, following this result, we expect that collecting more datasets would permit larger HSAs and higher EELS dose fractions, improving spectral signal-to-noise ratio (SNR).

### Proposed Workflow for Hollow Cryogenic Electron Ptychography with Spectroscopy

Based on our findings, we propose an integrated workflow for simultaneous structural and chemical analysis using hollow cryogenic electron ptychography and EELS, as illustrated in Fig. 4. The process consists of the following steps:

**Figure 4:**
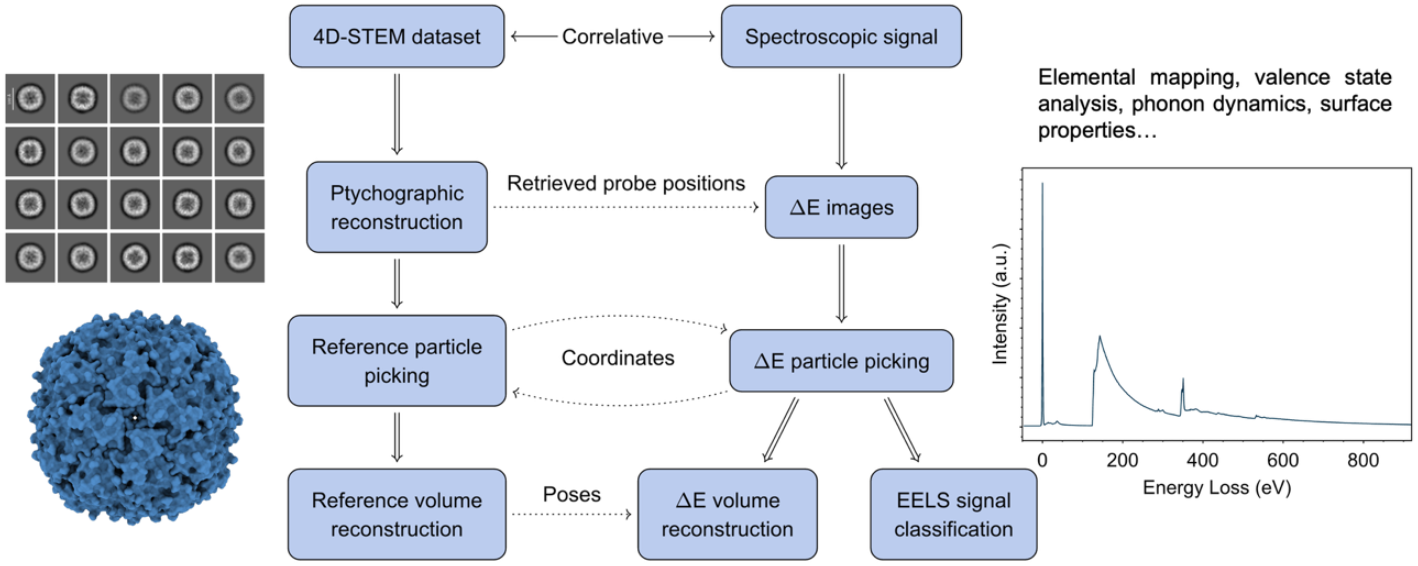
Schematic of the proposed workflow for hollow cryogenic electron ptychography with EELS. 4D-STEM data are acquired with a hollow detector that transmits the low-angle region of the BF disk to the EELS spectrometer, enabling simultaneous structural and spectral measurements. Hollow 4D-STEM data are reconstructed by ptychography to recover the phase and optionally probe positions, which are registered with the EELS spectrum at each scan position. The reconstructed phase images are processed by SPA to obtain a 3D density map. From the EELS spectra, ΔE images are extracted for all energy bins. Using the reference 3D map, particle coordinates and poses enable 3D ΔE mapping, yielding a 3D spatial map with an additional energy-loss dimension. Besides chemical mapping, the acquired EELS spectra can also be used for further chemical analysis. Illustrative EELS spectrum adopted from the EELS.info atlas^48^.

#### Data Acquisition

A 4D-STEM dataset is acquired using a hollow pixelated detector. The detector’s central opening passes through low-angle scattered electrons to an EELS spectrometer, while recording the remaining diffraction patterns. This enables simultaneous and co-registered acquisition of structural (diffraction, k_x_, k_y_) and spectral (EELS, ΔE) data at each scan position (s_x_, s_y_). When available, EDX measurements can be recorded in parallel for joint analysis.

#### 2D Ptychographic Reconstruction

The hollow 4D-STEM data (s_x_, s_y_, k_x_, k_y_) are processed using a ptychographic reconstruction algorithm to recover high-resolution phase images of the specimen.

#### 3D SPA Reconstruction

The reconstructed 2D phase images are imported into standard SPA software (e.g., cryoSPARC^32^, RELION^34,35^). Particles are picked, classified, and used for ab initio 3D reconstruction and refinement, yielding a high-resolution 3D density map and giving estimates to particle poses (coordinates, orientations).

#### Correlative Spectral Analysis

Since the EELS spectra and phase images are correlated at each probe position, the same particle coordinates and poses can be reused to classify spectra and improve their SNR. This is particularly valuable for beam-sensitive samples in the low-dose regime, where dose limitations yield noisy per-pixel spectra. Pose-based spectral classification enables averaging across similar poses, effectively increasing the dose per class and enabling more reliable extraction of chemical information, including elemental composition, bonding states, and oxidation states.

#### Energy-Resolved 3D Mapping

The correlated EELS spectra (pose, ΔE) can be used to generate energy-resolved 3D maps based on the reference high resolution structure from SPA. By reconstructing a 3D volume for each energy-loss channel, a 4D data cube (x, y, z, ΔE) can be created, directly correlating structural features with their chemical signatures, enabling a comprehensive analysis of the specimen within a single experiment.

## 3. Discussion

Simultaneous acquisition of spectral and structural information enables more accurate interpretation of macromolecular features and chemical states that would otherwise be ambiguous. We propose two instrument configurations with and without a hollow pixelated detector as shown in Fig. 1 Setup (a) and Setup (b), respectively. that can provide EELS alongside 4D-STEM.

In the Setup (a), a post-specimen deflector alternates the scattered electrons between a conventional pixelated detector and the EELS spectrometer. This configuration preserves complete diffraction patterns for conventional (non-hollow) ptychography and maximises the number of electrons for phase retrieval. However, the data acquisition is sequential, not truly simultaneous. To prevent unwanted exposure to the specimen during beam deflection between the detector and the EELS, a gun shutter may be required. The additional complexity and time delay introduced by the shutter and deflector make the experiment greater susceptible to drift and instabilities, resulting in degraded resolution and registration errors. Most importantly, the total dose budget increases due to dose partitioning across sequential modalities, making this approach potentially applicable for beam tolerant materials, but undesirable for beam-sensitive specimens, such as biological or organic materials.

In the Setup (b), the hollow detector configuration allows central part of the BF disk to passthrough to the EELS spectrometer while recording the remaining higher angle scattered electrons on the pixelated detector. This yields intrinsically simultaneous, perfectly co-registered structural and spectral datasets with a single total dose budget. For cryogenic biological applications prioritising dose efficiency, the hollow geometry is therefore the more practical solution provided the HSA is kept within empirically validated bounds.

The key trade-off in the hollow detector configuration is the systematic removal of low-angle diffraction information, producing HSA-dependent decline in ptychographic reconstruction quality and downstream SPA resolution. The dataset used for this analysis was obtained at an electron fluence of ∼35 e^-^/Å^2^, which is insufficient for high-SNR (fine-structure or trace-element) EELS on a per-particle basis; however, pose-based spectral averaging (using orientations and in-plane shifts recovered from the ptychographic SPA pipeline) can boost EELS SNR by aggregating equivalent views. Our results show that sub-nanometre 3D reconstructions are retained up to an HSA of 0.55α while allocating ∼30% of the dose to EELS. Under this setting, each particle can contribute ∼10 e^-^/Å^2^ to the spectrometer. To achieve single-atom EELS detection needs approximately 10^7^ e^-^/Å^2^, in order to reach such effective dose would require ∼10^6^ particles^14^. Data volumes of this scale are now common in conventional cryo-EM SPA as the total particle counts can routinely reach up to millions thanks to advances in detector technology and data collection automation^1,28,36,37^. This indicates that class-averaged EELS within the proposed hollow ptychography workflow is practical with contemporary scale experiments and should advance toward single-site sensitivity as particle counts and spectrometer efficiency improve.

Comparing to conventional STEM-HAADF with EELS, cryogenic electron ptychography with correlative EELS offers several advantages: (1) higher dose efficiency and robust phase recovery even at low dose^25,26^; (2) improved sensitivity to light elements that are commonly present in biomacromolecules but contribute weakly to HAADF contrast, due to high angle scattering weights heavier elements more favourably; (3) greater post-acquisition flexibility, since the 4D-STEM dataset is compatible with virtual imaging modes such as ADF and CoM, in addition to ptychography. These benefits make hollow ptychography a more versatile and effective tool for integrated structural and chemical analysis of biological specimens.

The correlation between ptychographic SPA with EELS spectra enables powerful analytical application. Ptychographic phase reconstructions are compatible with established SPA algorithms for particle picking, pose estimation, and 3D reconstruction and refinement. The recovered particle poses can be reapplied to the co-registered EELS data to perform pose-resolved spectral averaging and classification, increasing SNR and enabling orientation-dependent spectral analysis. Moreover, recent monochromated EELS can achieve energy resolutions of ≈4.2 meV at 30 kV and ≈8.0 meV at 100 kV^38,39^, sufficient to resolve vibrational modes that were previously unattainable in the very low loss regime. Vibrational EELS is highly sensitive to hydrogen^40-42^, allowing the detection and analysis of hydrogen bonding within organic compounds. This technique can be used for mapping of water distribution within biological specimens^43-45^, and potential tracking of labelled molecules via isotopes such as deuterium^46,47^. The core-loss region provides elemental composition (edge onsets) and local bonding, oxidation state, and electronic structure via near-edge fine structure^4^.

Furthermore, combining EELS with hollow cryogenic electron ptychography enables unified structural, chemical, and vibrational analysis. In principle, 3D density maps can be reconstructed for each energy-loss bin to generate a 4D (x, y, z, ΔE) dataset, directly correlating structural features with vibrational and chemical signatures, offering deeper insight into elemental composition, bonding environments, oxidation states, and ligand or solvent interactions in macromolecular assemblies. Besides, parallel EDX can be incorporated for complementary sensitivity to heavier elements. This framework extends cryogenic single-particle analysis from morphology alone to multimodal characterisation within a single acquisition.

## 4. Conclusion

In this work, we evaluated two configurations for integrating EELS with 4D-STEM: a beam-deflecting design and a hollow pixelated detector and their advantages and disadvantages. For beam-sensitive specimens, the hollow geometry is the more practical option because of its dose efficiency and the ability to perform simultaneous spectroscopy. Using a low-dose cryogenic apoferritin dataset, hollow ptychographic reconstructions maintained sub-nanometre 3D resolution up to 0.55α with ≈10^4^ particles while allocating ∼30% of the dose to EELS. Increasing particle numbers mitigated HSA-induced degradation, suggesting that larger datasets could support larger HSAs and higher EELS dose fractions, thereby improving spectral SNR and resolution. We proposed a workflow that couples 4D-STEM, ptychography, and SPA, leveraging pose-resolved averaging to boost spectral SNR and enabling energy-resolved 3D mapping co-registered with the structural reconstruction. Current limitations arise from reconstruction algorithm maturity and pipeline integration. Further developments in ptychographic reconstruction algorithm, detector geometry, and tighter integration with SPA, coupled with advances in EELS instrumentation such as monochromators and aberration correctors, should improve robustness, dose efficiency, and attainable resolution of this technique, enabling comprehensive, simultaneous structural–chemical characterisation of biological macromolecules.

## Acknowledgements

P.W. acknowledges funding from BBSRC International Institutional Partnership Fund and the University of Warwick Research Development Fund (RDF) 2021-22 Science Development Award. X.W. and Y.L. are supported by studentships within the Engineering and Physical Sciences Research Council supported Centre for Doctoral Training in Modelling of Heterogeneous Systems, under Grant Nos. EP/Y035429/1 and EP/S022848/1, respectively.

## Author contributions

P.W. conceived the overall project and supervised X.W. and Y.L., who performed ptychographic reconstructions, single particle analysis and related simulations. X.W. and P.W. wrote the manuscript. All authors discussed the results and contributed to the final version of the paper.

## Data availability

All data that support this study will be posted on a public website upon publication.

## Code availability

The MultiHollowPtycho source code is available at the GitHub repository:

## Supplementary Information

**Supplementary Figure 1:**
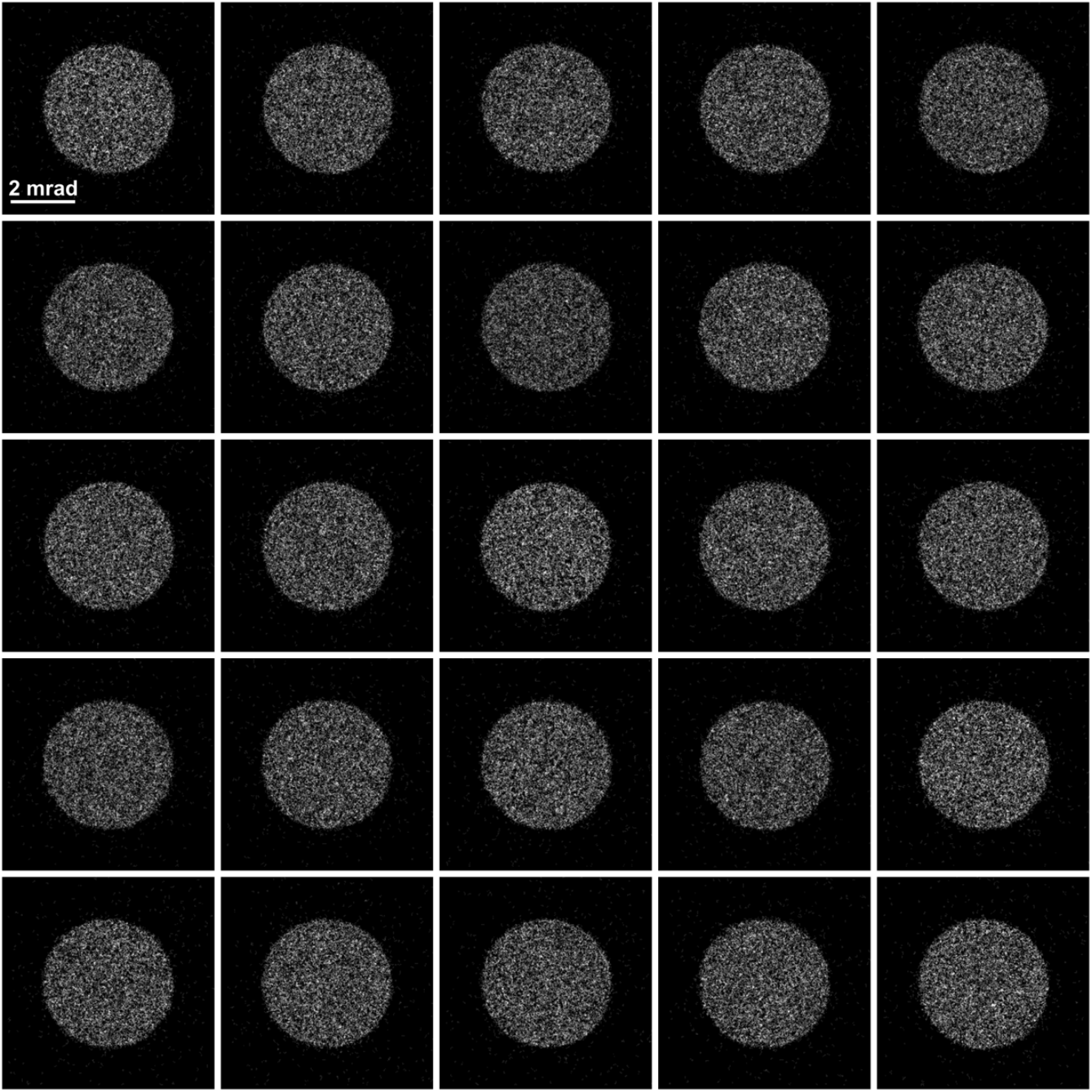
Diffraction patterns in a 5 × 5 array taken from the full 128 × 128 dataset, each diffraction pattern has 256 × 256 pixels.

**Supplementary Figure 2:**
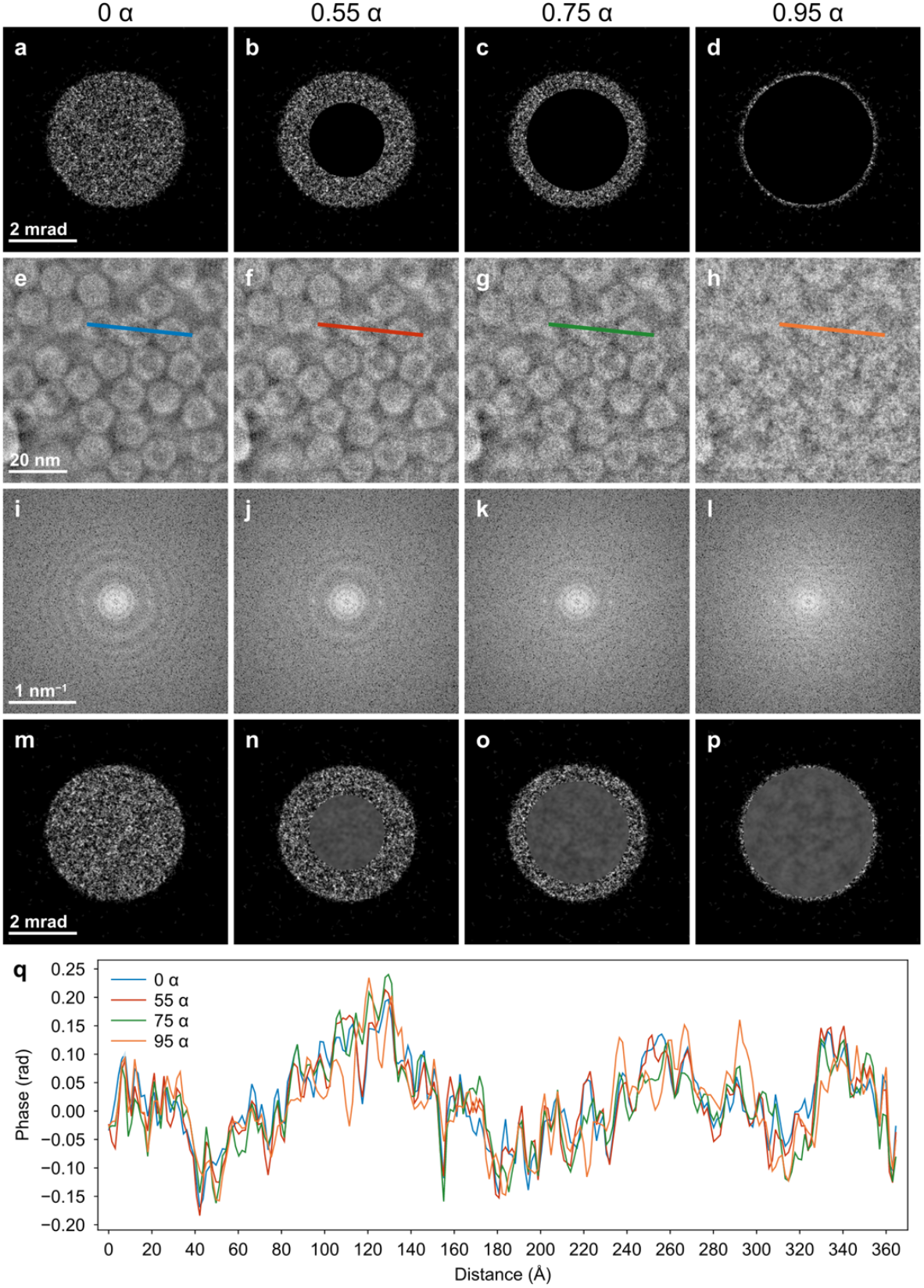
Ptychographic reconstruction of apoferritin using different HSAs. **a**-**d** Diffraction patterns with HSA values of 0, 0.55, 0.75 and 0.95× of the probe CSA (α). **E**-**f** Reconstructed phase images of the corresponding HSAs. **i**-**l** Power spectra of the corresponding phase images. **m**-**p** Reconstructed diffraction patterns at the corresponding HSA. **q** Line profiles of the reconstructed phase images as indicated in **e**-**h**.

**Supplementary Figure 3:**
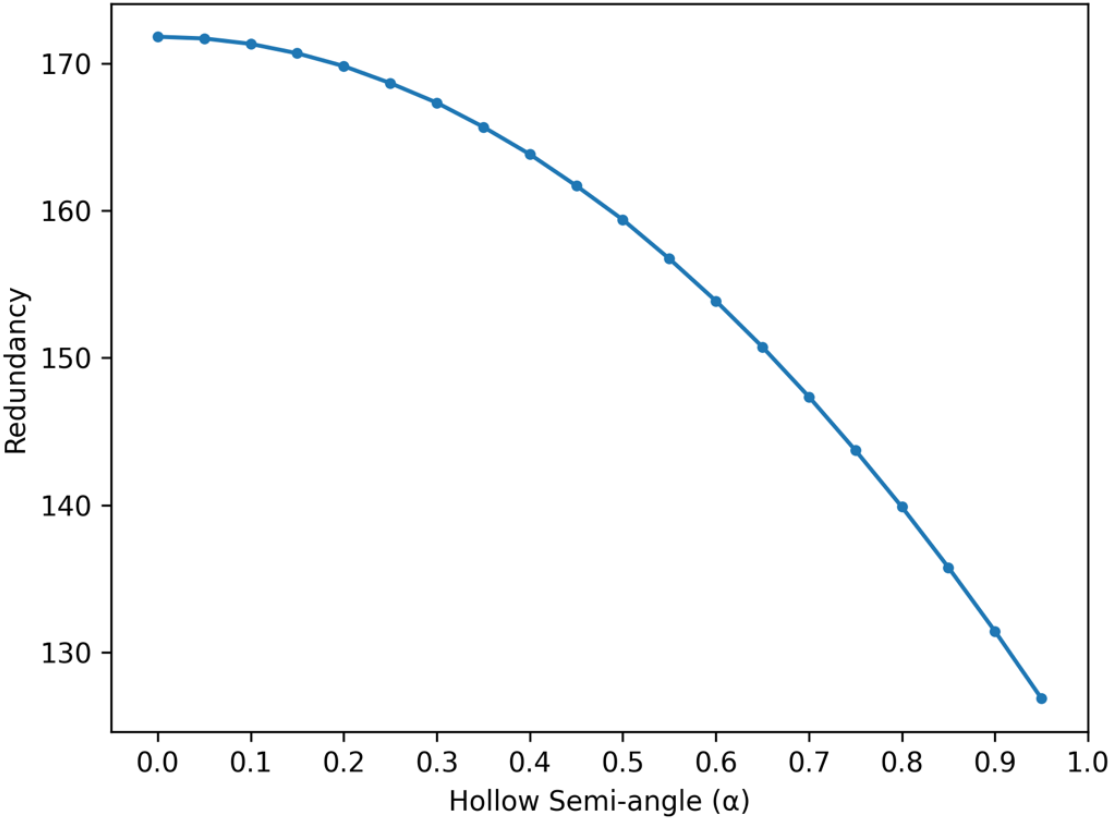
Dataset redundancy for ptychographic reconstructions as the HSA increases.

**Supplementary Figure 4:**
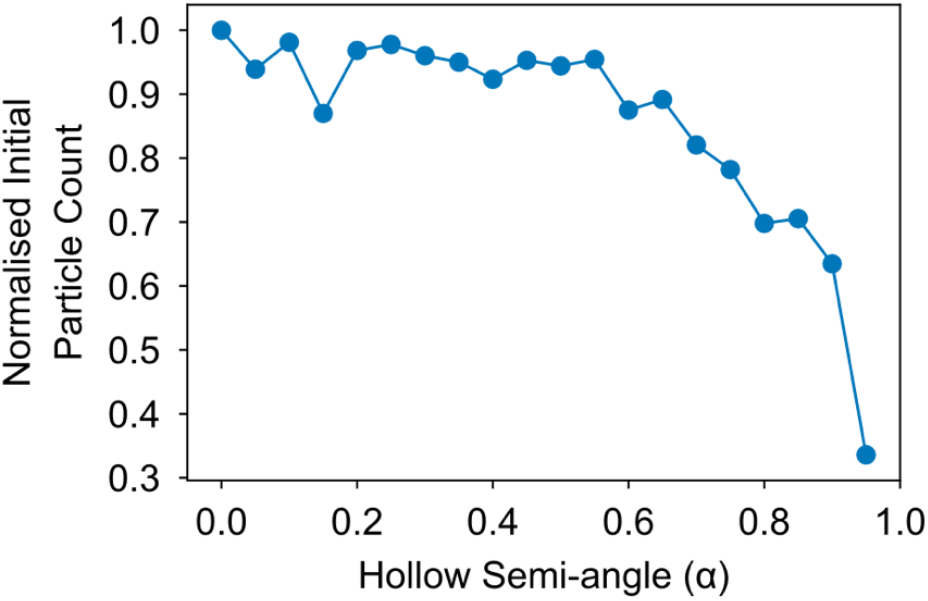
Ratio of initially selected particles at each hollow semi-angle compared to the number of particles picked from the reference reconstruction using the full diffraction patterns.

**Supplementary Figure 5:**
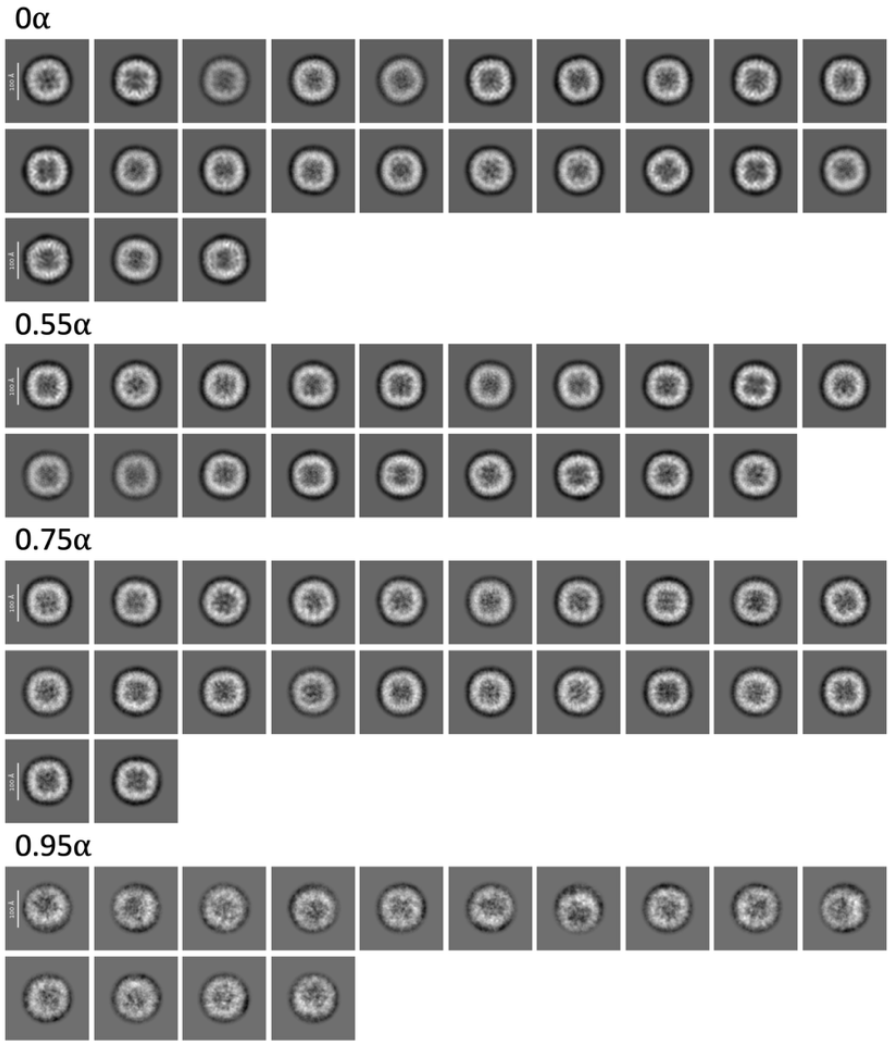
2D class averages of apoferritin images used for *ab initio* reconstructions.

**Supplementary Figure 6:**
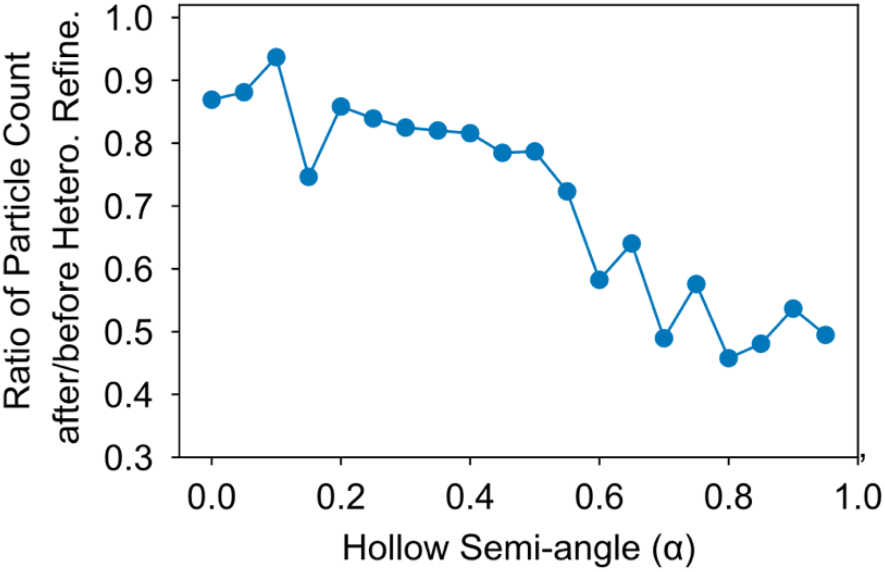
Ratio of initially selected particles assigned to the best heterogeneous class after *ab initio* reconstruction, representing the effective usable particle yield.

**Supplementary figure 7:**
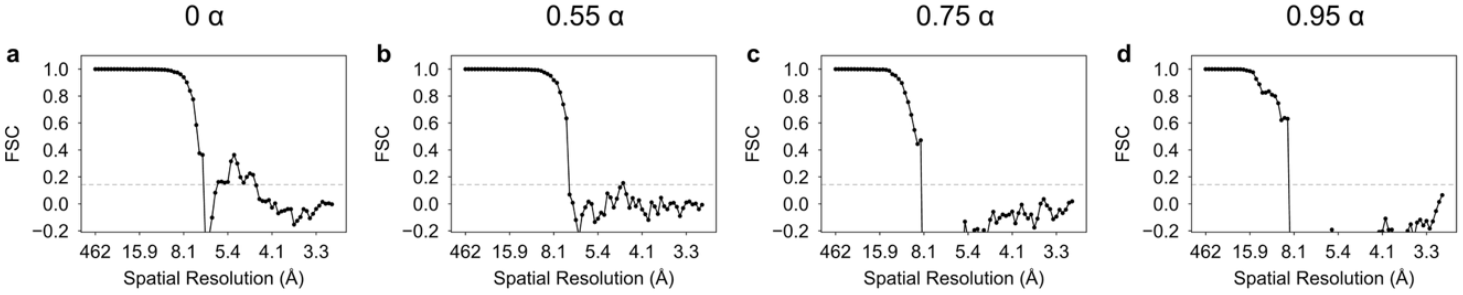
FSC curves for apoferritin reconstructions using all the available particles at different HSA.

**Supplementary Figure 8:**
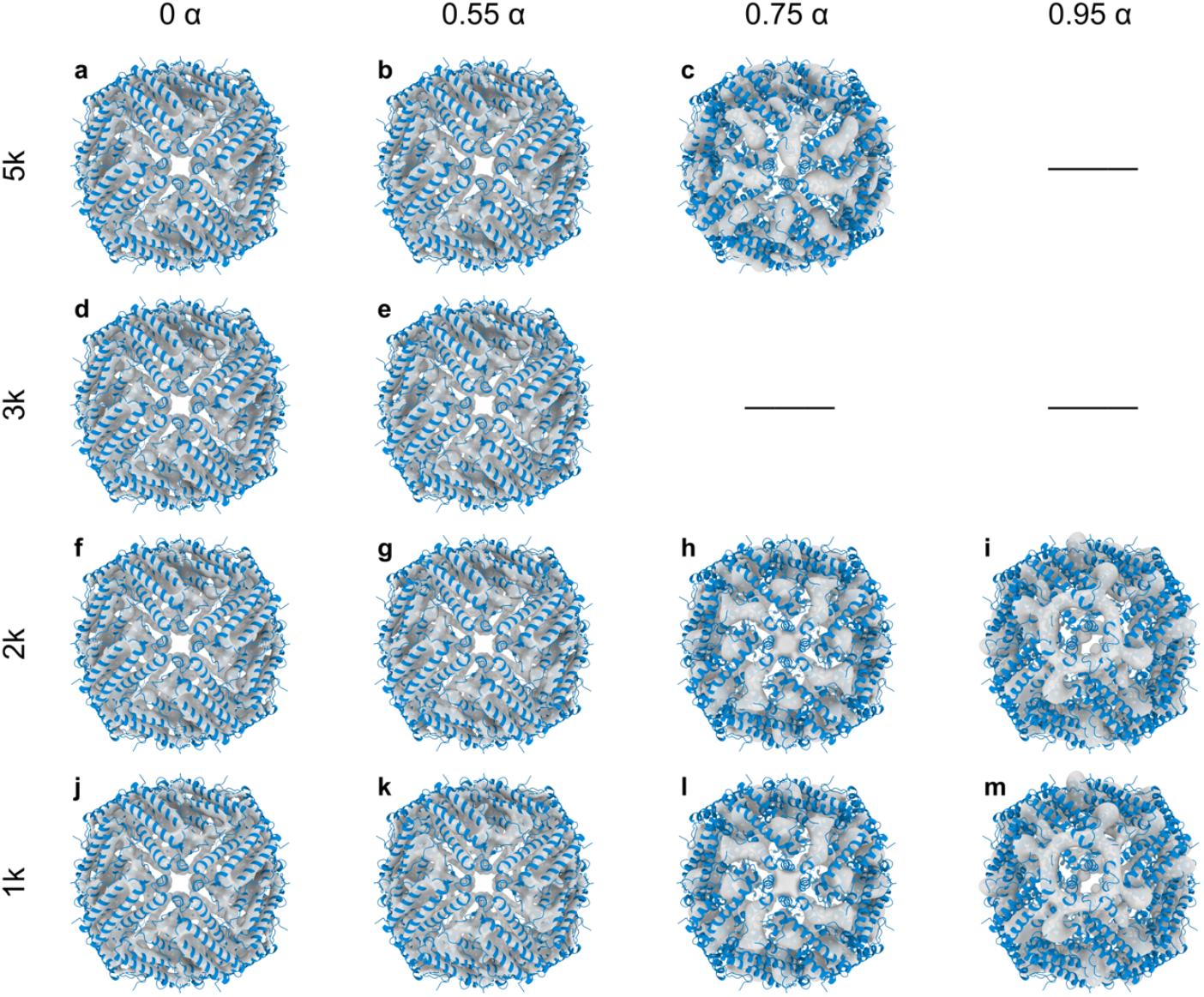
3D reconstructions of the apoferritin protein with different HSA and available particle counts. **a**–**c** Maps from datasets at HSA = 0, 0.55, and 0.75 × of the probe CSA (α) using 5,000 particles. **d**-**e** Maps using 3,000 particles at the corresponding HSAs. **f**–**i** Maps using 2,000 particles at 0α, 0.55α, 0.75α and 0.95α. **j**–**m** Maps using 1,000 particles across HSAs. Note: particle count refers to those used for the final local refinement and were randomly sampled from the best 3D class. For the 5,000-particle condition, all available particles from the dataset were used for *ab initio* reconstruction; for smaller reconstructions, 5,000 particles were used at the *ab initio* stage.

**Supplementary figure 9:**
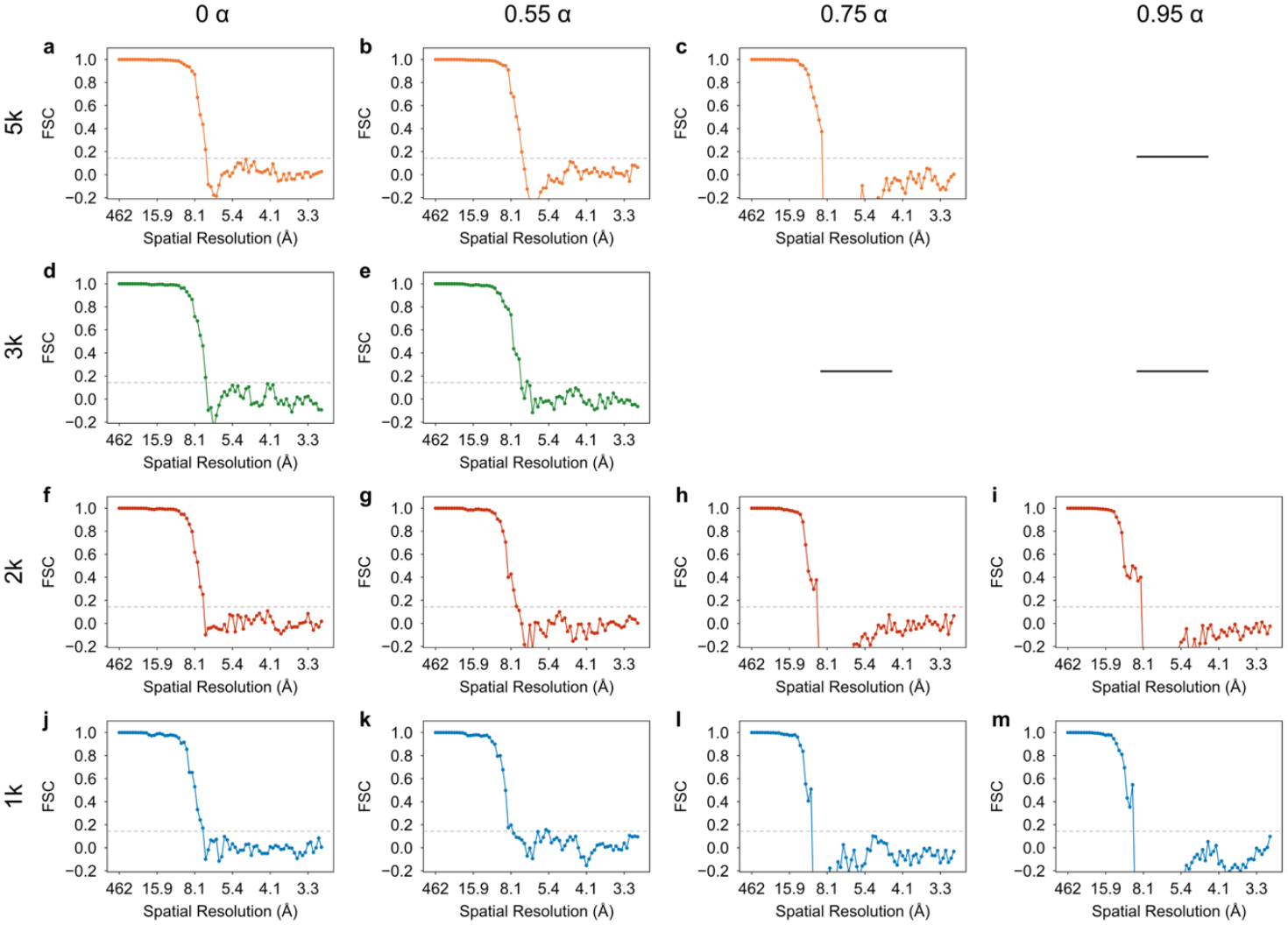
FSC curves for apoferritin reconstructions with different HSA and particle count. **a**–**c** FSCs for maps reconstructed from datasets at HSA = 0, 0.55 and 0.75 × of the probe CSA (α) using 5,000 particles. **d**–**e** FSCs using 3,000 particles at the corresponding HSAs. **f**–**i** FSCs using 2,000 particles at 0α, 0.55α, 0.75αand 0.95α. **j**–**m** FSCs using 1,000 particles at the corresponding HSAs. Note: particle counts denote those used for the final local refinement and were randomly sampled from the best 3D class. For the 5,000-particle condition, all available particles from the dataset were used for *ab initio* reconstruction; for smaller reconstructions, 5,000 particles were used at the *ab initio* stage.

**Supplementary Table 1:** Parameters for cryogenic electron ptychographic data acquisition^1^ and hollow reconstruction.

| Dataset | Setting (1) | Setting (2) | Setting (3) | Setting (4) |
| --- | --- | --- | --- | --- |
| Accelerating voltage (kV) | 300 | 300 | 300 | 300 |
| Convergence Semi-angle (mrad) | 4.0 | 4.0 | 4.0 | 4.0 |
| Maximum detection semi-angle (mrad) | 6.58 | 6.58 | 6.58 | 6.58 |
| Sampling in real space (Å) | 1.5 | 1.5 | 1.5 | 1.5 |
| Defocus (μm) | 2 | 2 | 2 | 2 |
| Probe size (nm) | 16 | 16 | 16 | 16 |
| Raster scan dimension | 128 × 128 | 128 × 128 | 128 × 128 | 128 × 128 |
| Step size (nm) | 2 | 2 | 2 | 2 |
| Diffraction Pattern dimension (pixels) | 256 × 256 | 256 × 256 | 256 × 256 | 256 × 256 |
| Dwell time per diffraction pattern (μs) | 250 | 250 | 250 | 250 |
| Dose (e<sup>-</sup>/Å<sup>2</sup>) | 35 | 35 | 35 | 35 |
| Hollow semi-angle (mrad) | 0.0 | 2.2 | 3.0 | 3.8 |
| Overlap ratio | 0.84 | 0.84 | 0.84 | 0.84 |
| Redundancy $\sigma_{\text{pty}}$ | 172 | 157 | 144 | 127 |

### Supplementary Note 1: Experimental 4D-STEM dataset of apoferritin

The cryogenic 4D-STEM datasets of apoferritin, previously reported by Küçükoğlu et al^1^ was used in this study. Apoferritin were applied to UltraFoil gold 1.2/1.3 grids and plunge-frozen in liquid ethane using a Leica EM GP2.

The 4D-STEM dataset was acquired on a probe-corrected Thermo Fisher Scientific Titan Krios G4 with a spherical aberration corrector and a cold field emission gun at an accelerating voltage of 300 kV. The microscope was operating at 4 mrad and a defocus of -2 µm. The 4D-STEM data was recorded using half of the Dectris ELA hybrid pixel detector (256 × 256 pixels) with a dwell time of 250 µs, a probe size of ∼16 nm, a step size of 2 nm and a scanning grid of 128 × 128. The average total electron dose applied to the sample is ∼35 e^−^/Å^2^.

For further details about the sample preparation and experimental setup, please refer to reference^1^.

### Supplementary Note 2: Reconstruction algorithm for hollow ptychography

The reconstruction method used in this work is based on a modified version of py4DSTEM’s batched gradient descent implementation for single slice ptychography^2^. The ptychographic phase retrieval problem can be described by Eq. (1-3), where the exit wave Ψ_i_ (***r***) at scan position *i* is the product of the corresponding probe *P*(***r***) and object 𝒪(***r*** − ***R***_***i***_).

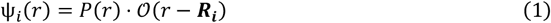

The far-field diffraction pattern is the Fourier transform of the exit wave:

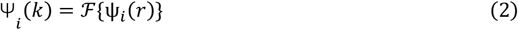

Which is constrained by the experimentally measured intensity *I*_*i*_ (***k***):

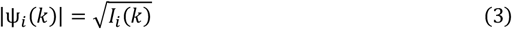

Given an initial estimation of the probe and object function, the forward exit wave function can be computed and updated by projecting the Fourier constraint:

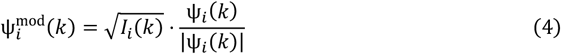

At this step a mask matching the shape of the hollow detector is applied, to exclude the missing regions in the diffraction pattern, such that the retrieved signal can still be utilized for updating both the probe and the object. The difference between the modified exit wave and the current estimate:

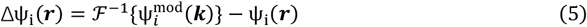

In each iteration, the algorithm would randomly permute all probe positions and subdivide them into mini batches, each mini batch *B* is of the size *M* ≪ *N*, where *N* is the total number of diffraction patterns.

For each mini batch, the object function could be updated as:

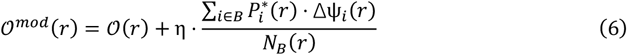

Here *η* is the step size that controls the convergence rate, and *N*_*B*_ (***r***) is the probe normalization factor over the current batch:

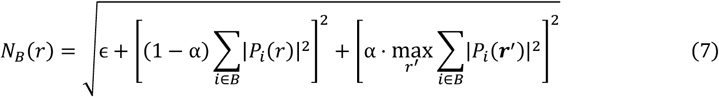

Here *ϵ* is a small constant added for numerical stability, and *α* is the normalization minimum parameter.

Similarly, the probe can be updated by:

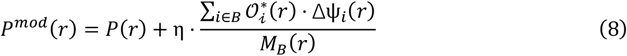

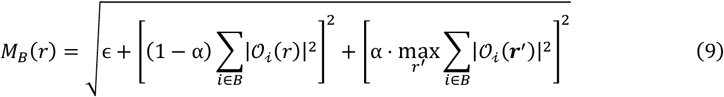

## Supplementary Note 3: Single particle reconstruction procedures in cryoSPARC

Reconstructed ptychographic phase images were saved as MRC file and then imported into CryoSPARC for SPA^3^. Initial particle picking was performed using a blob picker, followed by 2D classification to generate templates for subsequent automated picking. Several rounds of iterative 2D classification were performed to curate the particle stack, using 200, 100, 50, and finally 25 classes, with each classification job running for 40 online-EM iterations. Particles were initially extracted into a 128-pixel box, then re-extracted into a 154-pixel box for the final classification rounds before moving into 3D reconstruction.

The curated particle set was used for ab-initio reconstruction based on a stochastic gradient descent algorithm with octahedral symmetry imposed and a class similarity of 0.9. The resulting 3D map was subjected to homogeneous, non-uniform, and local refinement with re-centering, followed by particle re-extraction. A two-class heterogeneous refinement was performed with an initial low pass filter of 15 Å to isolate the best-resolved particle stack, which was then subjected to a final local refinement. The resolution of the final density maps was estimated using the gsFSC with a 0.143 cutoff^4^.

## Supplementary Note 4: Dataset redundancy evaluation

To quantify the degree of data redundancy in a ptychographic dataset, the metric *σ*_*pty*_ can be computed as^5^:

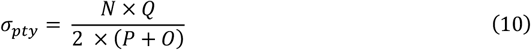

Where *N* is the number of diffraction patterns used in the reconstruction, *Q* is the number of pixels there are in each diffraction pattern recorded on a hollow detector, *P* is the total number of pixels in the probe function and *O* is the total number of pixels in the reconstructed object function.

The calculated data redundancy *σ*_*pty*_ for the dataset used can be found in Supp. Table 1. The *σ*_*pty*_ value at each HSA is shown in Supp. Fig. 3.

